# LAIR1 prevents excess inflammatory tissue damage in *S. aureus* skin infection and Cutaneous T-cell Lymphoma

**DOI:** 10.1101/2024.06.13.598864

**Authors:** Hannah K. Dorando, Evan C. Mutic, Kelly L. Tomaszewski, Ling Tian, Mellisa K. Stefanov, Chaz C. Quinn, Deborah J. Veis, Juliane Bubeck Wardenburg, Amy C. Musiek, Neha Mehta-Shah, Jacqueline E. Payton

## Abstract

Patients with cutaneous T cell lymphoma (CTCL) experience high morbidity and mortality due to *S. aureus* skin infections and sepsis, but the causative immune defect is unclear. We previously identified high levels of LAIR2, a decoy protein for the inhibitory receptor LAIR1, in advanced CTCL. Mice do not have a LAIR2 homolog, so we used *Lair1* knock-out (KO) mice to model LAIR2 overexpression. In a model of subcutaneous *S. aureus* skin infection, *Lair1* KO mice had significantly larger abscesses and areas of dermonecrosis compared to WT. *Lair1* KO exhibited a pattern of increased inflammatory responses in infection and sterile immune stimulation, including increased production of proinflammatory cytokines and myeloid chemokines, neutrophil ROS, and collagen/ECM remodeling pathways. Notably, *Lair1* KO infected skin had a similar bacterial burden and neutrophils and monocytes had equivalent *S. aureus* phagocytosis compared to WT. These findings support a model in which lack of LAIR1 signaling causes an excessive inflammatory response that does not improve infection control. CTCL skin lesions harbored similar patterns of increased expression in cytokine and collagen/ECM remodeling pathways, suggesting that high levels of LAIR2 in CTCL recapitulates *Lair1* KO, causing inflammatory tissue damage and compromising host defense against *S. aureus* infection.

## INTRODUCTION

Cutaneous T-cell Lymphoma, or CTCL, is a cancer of skin-homing T-lymphocytes. Patients with late-stage disease have poor survival rates, and the only curative treatment is allogeneic stem cell transplant. Advanced CTCL causes inflammatory tissue damage in CTCL-involved skin and defects in cell-mediated immunity that lead to frequent *Staphylococcus aureus* (*S. aureus*) skin and soft tissue infections (SSTI) associated with high morbidity and mortality (1–3). Notably, the mechanisms that lead to defects in cell-mediated immunity and inflammatory tissue damage in CTCL remain undefined (4–7).

We previously identified aberrant epigenetic regulation and high expression of leukocyte-associated immunoglobulin-like receptor 2 (LAIR2) in advanced and treatment-resistant CTCL (8). LAIR2 and its paralog, LAIR1, are Ig-like receptors that bind collagens and other proteins with collagen-like domains, including complement pathway proteins such as C1q, which are important in host defense and tissue remodeling (9–12). LAIR1 is a transmembrane inhibitory receptor with two intracellular ITIM motifs. LAIR2, a secreted protein nearly identical to the extracellular domain of LAIR1, acts as a decoy receptor, reducing LAIR1 inhibitory signaling and thereby increasing the level of activation signals in the cell (10). We and others demonstrated that LAIR1 is highly expressed on macrophages and other innate immune cells; in contrast, LAIR2 expression is largely restricted to T and NK cells (13–15). Upon binding collagen domain-containing ligands, LAIR1 signaling via SHP1 and SHP2 phosphatases negatively regulates many cellular immune functions, including cytokine production, calcium signaling, and differentiation (16–20).

Available evidence supports the notion that LAIR proteins likely function in inflammatory responses in infection and inflammatory diseases. Indeed, we and others demonstrated that LAIR1 and LAIR2 expression varies in cell type– and factor-specific patterns in response to pathogen– and host-derived inflammatory factors (13–15). Decreased levels of LAIR1 are associated with rheumatoid arthritis, systemic lupus erythematosus, inflammatory bowel disease, and psoriasis (21, 22). LAIR1 expression in innate immune cells (monocytes, NK cells, neutrophils) may play a modulatory role in immune response to RSV infection (21, 23, 24), severe COVID-19 (25), and pediatric malarial infection (26, 27). These reports support a role for LAIR proteins in inflammatory diseases and immune response to viral and malarial infection, but the underlying mechanisms remain to be defined and a role in bacterial infection has not been tested to our knowledge.

We previously identified LAIR2 upregulation in CTCL patients, a population with increased susceptibility to invasive *S. aureus* SSTI (1–3). Therefore, we sought to define the role of LAIR proteins in a mouse model of *S. aureus* SSTI. Because mice do not have a gene to produce LAIR2, *Lair1* KO mice provide a simple and powerful way to achieve a murine model for loss of LAIR1 inhibitory signaling (10, 28). We used a well-characterized mouse model of *S. aureus* SSTI established by the Bubeck Wardenburg lab in which mice are subcutaneously inoculated with a high dose of a clinically relevant isolate of methicillin-resistant *S. aureus* (MRSA), USA300/LAC (29), causing areas of abscess and dermonecrosis (death of overlying epithelium) that spontaneously resolve within 2 weeks in wild type mice (30–32).

In the results of *in vivo* and *in vitro* studies presented here, we discovered that loss of LAIR1 causes striking differences in the response to *S. aureus*. In skin infections, *Lair1* KO mice developed much larger areas of abscess and dermonecrosis that did not resolve within the experimental timeframe, with no difference in bacterial burden. *Lair1* KO skin lesions also had higher levels of pro-inflammatory cytokines; these same cytokines were elevated in *Lair1* KO bone marrow-derived macrophages (BMDM) infected with MRSA *in vitro*. *Lair1* KO neutrophils produced higher levels of reactive oxygen species (ROS), yet demonstrated equivalent phagocytosis of *S. aureus* compared to WT. In CTCL skin biopsies from our studies and publicly available datasets, several of the same cytokines were elevated in advanced disease, *notably without infection*. In addition, we found that *S. aureus* skin lesions from *Lair1* KO exhibited increased expression of genes in extracellular matrix remodeling, collagen biosynthesis, and collagen degradation pathways. The same collagen/ECM pathways were upregulated in TLR2-activated BMDMs from *Lair1* KO and several of the same collagen and ECM genes were also upregulated in CTCL skin biopsies with advanced disease. Taken together, our findings reveal a novel role for LAIR1 in bacterial infection and support the notion that LAIR1 inhibitory signaling in macrophages modulates immune response during *S. aureus* infection and prevents excess tissue damage. These studies also provide insight into a potential mechanism for involvement of LAIR2 in immune dysfunction as well as inflammatory skin and tissue damage in CTCL.

## RESULTS

### LAIR2 is elevated in CTCL tumor cells

We previously reported that high levels of LAIR2 are associated with advanced and therapy-resistant cutaneous T-cell lymphoma (CTCL), a disease marked by inflammatory damage to skin and underlying soft tissue, as well as deficits in cellular immunity that cause frequent and severe *S. aureus* skin and soft tissue infections (SSTI) (1–3, 8). As an inhibitory Ig-like receptor, LAIR1 signaling counteracts inflammatory signaling in immune cells; LAIR2 acts as a decoy receptor and disrupts LAIR1 inhibitory signals (10). Therefore, we hypothesized that reduced LAIR1 inhibitory signaling promotes inflammation in CTCL and alters immune response to *S. aureus* infection.

To begin to test this hypothesis, we first examined single cell (sc)RNA-seq from 16 serial peripheral blood samples from CTCL patients. CD4+ cells were enriched from peripheral blood mononuclear cells (PBMCs) to enhance capture of CTCL cells, which are CD4+ in the vast majority of patients (33). We used 10x Genomics Chromium Single Cell 5’ Gene Expression and V(D)J Immune Profiling (v1.0) to generate expression and TCR (T-cell receptor) repertoire profiles. In total, 92,496 cells passed QC filters; annotation of cell type was performed using the Human Primary Cell Atlas (HPCA) data set as reference (34). Of 92,496 cells from all samples, the majority were CD4+ T cells (76,924); monocytes, which are also CD4+, comprised most of the remaining cells (**Fig. 1A** and **S1A**). Automated cell annotations were confirmed using canonical marker genes: pan T-cell – *CD3D*, T cell receptor beta (*TRBC2*); helper (*CD4*) versus cytotoxic (*CD8A*, *CD8B*); CTCL – *TOX*, *KIR3DL2*, *PLS3*; and monocytes – *CD14*, *LYZ*, *S100A9* (35, 36) (**Fig. S1B-D**). Cell type assignments show a clear separation of CD4+ non-T-cells, most of which are monocytes, and T cells (**Fig. 1A**). In accordance with our previous work (8, 13), we found that LAIR1 was expressed primarily in myeloid cells (**Fig. 1B**), while LAIR2 is expressed highly in CTCL T cells (**Fig. 1C**). Thus, we hypothesized that LAIR2 secreted from CTCL cells blocks LAIR1 inhibitory signaling in myeloid cells, causing increased inflammation and associated tissue damage, which negatively impact host defense and create a permissive environment for invasive *S. aureus* infection (**Fig. 1D**).

**Figure 1.**
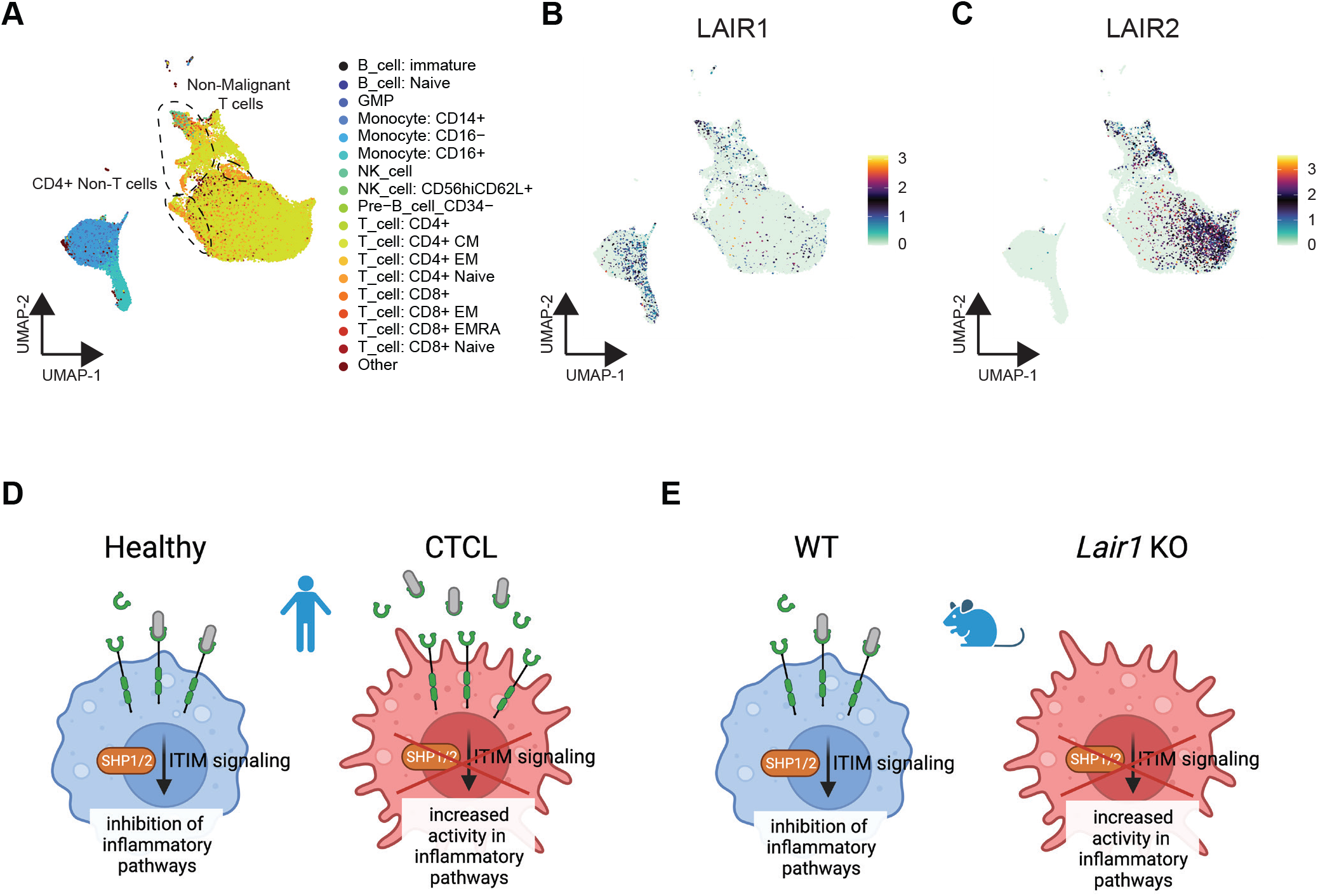
LAIR2 is overexpressed in CTCL and reduces LAIR1 signaling, analogous to *Lair1* KO. (**A** – **C**) CD4+ cells were isolated from CTCL patient peripheral blood and analyzed by single cell RNA-seq. UMAP projections show 92,496 mononuclear cells from 16 peripheral blood samples from 6 CTCL patients across multiple time points. (**A**) Cell type assignment by highest spearman rho of purified immune populations in the Human Primary Cell Atlas. UMAP plots show expression of LAIR1 (**B**) and LAIR2 (**C**). (**D**) Cartoon depicts hypothesis that LAIR2 overexpression reduces LAIR1 signaling in CTCL. (**E**) Cartoon depicts mouse model for loss of LAIR1 signaling via KO of *Lair1*.

To test the role of LAIR proteins in *S. aureus* skin infection, we used *Lair1* knock-out (KO) mice to achieve loss of LAIR1 signaling because mice do not have a *LAIR2* homolog (**Fig. 1E**) (10, 28).

### LAIR1 is protective in S. aureus skin infection

We employed a mouse model of *S. aureus* SSTI in which wild-type (WT) and *Lair1* KO mice were inoculated subcutaneously with 1×10^7^ CFU *S. aureu*s USA300/LAC. In WT mice, this results in a self-limited skin infection, consisting of abscess and dermonecrosis, that resolves spontaneously over the course of two weeks (30–32). The infectious lesions consist of an abscess, which has a collagen-rich barrier created to contain the bacteria, leukocytes, and debris generated within 1 day post-infection (dpi). Dermonecrosis is defined as death of epithelium overlying the abscess. In WT mice, dermonecrosis develops at 2 dpi. Strikingly, *Lair1* KO mice develop abscesses that are two-fold larger on average (**Fig. 2A**) and develop areas of dermonecrosis that are two-fold larger and develop more rapidly (by 1dpi, **Fig. 2B**) compared to WT mice. Differences in abscess and dermonecrosis sizes were seen beginning at 1 dpi and persisted throughout the experimental timeline, with the greatest differences at 1-8 dpi. This phenotypic difference early in infection suggests a predominant role for myeloid cells. Additionally, while WT mice completely resolve the infection by 14 dpi, an average of over 10% of lesion area remained in *Lair1* KO at 14 dpi. Surprisingly, *S. aureus* CFU recovered per skin lesion biopsy at 2 dpi was no different in WT compared to *Lair1* KO (**Fig. 2C**). *S. aureus* CFU recovered from spleen at 2 dpi and 5 dpi was not statistically different in *Lair1* KO compared to WT (**Fig. 2D**). Representative hematoxylin and eosin (H&E) staining of tissue excised from mice with SSTI at 2 dpi demonstrates the larger areas of abscess and dermonecrosis in *Lair1* KO compared to WT (**Fig. 2E**). Together, these findings suggest that absence of LAIR1 in *S. aureus* SSTI causes greater tissue damage yet does not meaningfully impact bacterial clearance.

**Figure 2.**
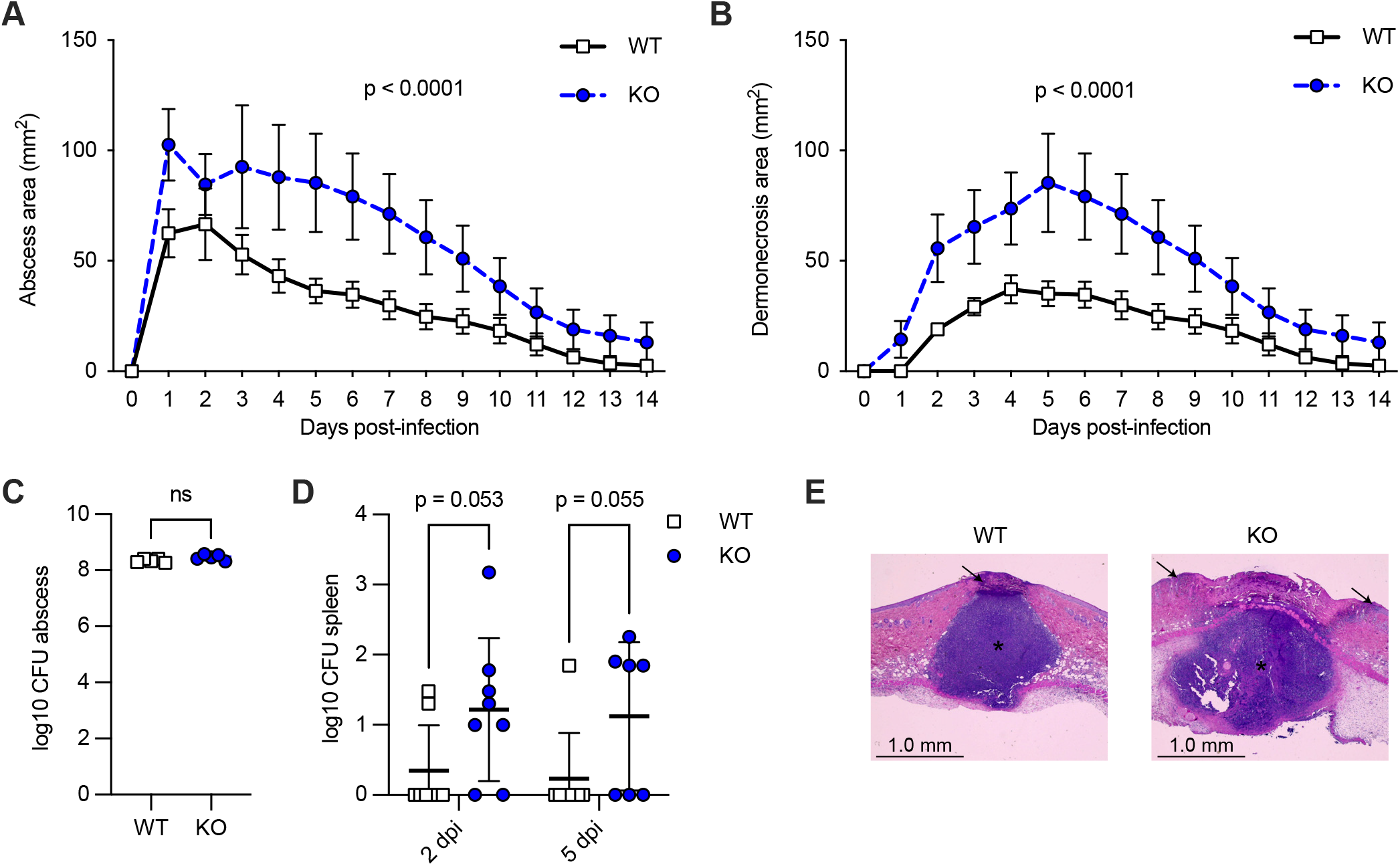
LAIR1 is protective in *S. aureus* skin infection. WT and *Lair1* KO mice were infected subcutaneously with 1×10^7^ CFU *S. aureus* USA300 LAC. Lesions were monitored over 14 days and quantified as abscess (**A**) and dermonecrosis (**B**) area. (noninvasive measurements, area calculated: A = ([pi/2] x l x w). Repeated measures two-way ANOVA with Bonferroni post-hoc tests. Results are representative of five independent experiments (n = 40 WT, n = 40 *Lair1* KO). Bacterial growth was quantified as CFU recovered from homogenized skin punch biopsies collected at 2 days post-infection (dpi) (**C**) and from spleen at 2 dpi and 5 dpi (**D**) (unpaired t-test). (**E**) Representative H&E stains of lesions at 2 dpi. Abscesses are indicated by stars and arrows indicate areas of dermonecrosis, which are larger in *Lair1* KO.

α-Hemolysin, or Hla, is a pore-forming *S. aureus* virulence factor that causes lysis of host cells. Isogenic *hla*-negative *S. aureus* strains caused significantly smaller skin lesions with little to no dermonecrosis compared to WT strains (32, 37). To determine whether the larger abscess and dermonecrosis areas in *Lair1* KO SSTI were dependent on Hla-mediated tissue damage, we used an isogenic *hla*-negative strain of USA300/LAC (Δhla)(37). We inoculated each WT and *Lair1* KO mouse with USA300/LAC on one flank and Δ*hla* USA300/LAC on the opposite flank and compared abscess area (**Fig. S2A-B**) and dermonecrosis area (**Fig. S2C-D**) over two weeks. Overall, infection with the Δ*hla* strain reduced the abscess and dermonecrosis sizes in both WT and *Lair1* KO compared to the wild-type strain. However, Δ*hla* abscesses in *Lair1* KO were larger than WT mice with wild-type or Δ*hla* infections (**Fig. S2A**). Moreover, areas of dermonecrosis in Δ*hla*-infected *Lair1* KO were similar to wild-type infections in WT mice and were larger than Δ*hla* infections in WT mice (**Fig. S2C**).

Statistical analysis of these results confirmed that there are significant differences due to *S. aureus* strain and genotype. A three-way ANOVA was conducted to compare the main effects of genotype and *S. aureus* strain as well as their interaction effects on either size of abscess (**Fig. S2B**) or dermonecrosis (**Fig. S2D**). Effect of genotype (WT vs. *Lair1* KO) was statistically significant for both abscess and dermonecrosis at p < 0.0001. Effect of *S. aureus* strain (USA300 vs. Δ*hla*) was statistically significant for both abscess (p = 0.0008) and dermonecrosis (p < 0.0001). The two-way interaction effect for genotype and *S. aureus* strain was not significant for abscess, but was significant for dermonecrosis at p = 0.0249, indicating that the impact of genotype on dermonecrosis depended on *S. aureus* strain. To determine significant differences, two-way ANOVAs with Bonferroni adjustment for alpha significance level were used as post-hoc tests to determine significant differences within the dermonecrosis two-way interaction effect: dermonecrosis is significantly greater in KO than WT when using USA300 strain (p = 0.0004) and showed a non-significant trend toward a larger size in KO versus WT when using Δ*hla* strain (p = 0.0692). This result was expected due to the overall low amount of dermonecrosis in Δhla infection. Taken together, these data demonstrate that the increased abscess and dermonecrosis sizes in *S. aureus*-infected *Lair1* KO mice were not solely due to the activity of Hla. Therefore, we next sought to define host-derived differences in response to *S. aureus* infection.

### Lair1 KO mice produce higher levels of pro-inflammatory cytokines in skin lesions

Because we observed larger lesions and more tissue damage in *S. aureus* SSTI in *Lair1* KO with no difference in bacterial burden, we hypothesized that *Lair1* KO mice produce increased pro-inflammatory cytokines. We therefore measured production of 32 cytokines and growth factors from skin punch biopsies from WT and *Lair1* KO SSTI at 18 hours post-infection by multiplex cytokine array. *Lair1* KO skin had an overall pattern of increased pro-inflammatory cytokine expression (**Fig. 3A**), with five cytokines exhibiting significantly higher levels compared to WT (**Fig. 3B-F**). Neutrophil-activating chemokines CXCL1 and CXCL2 are both significantly increased, with CXCL1 elevated by more than 100% and CXCL2 elevated by about 8% on average (**Fig. 3B-C**). Monocyte-activating chemokines CCL2 (MCP-1) and CCL3 (MIP-1α) were both elevated by greater than 60% on average (**Fig. 3D-E**). IL-1β, which is classically associated with response to bacterial infection, was increased by over 90% on average (**Fig. 3F**).

**Figure 3.**
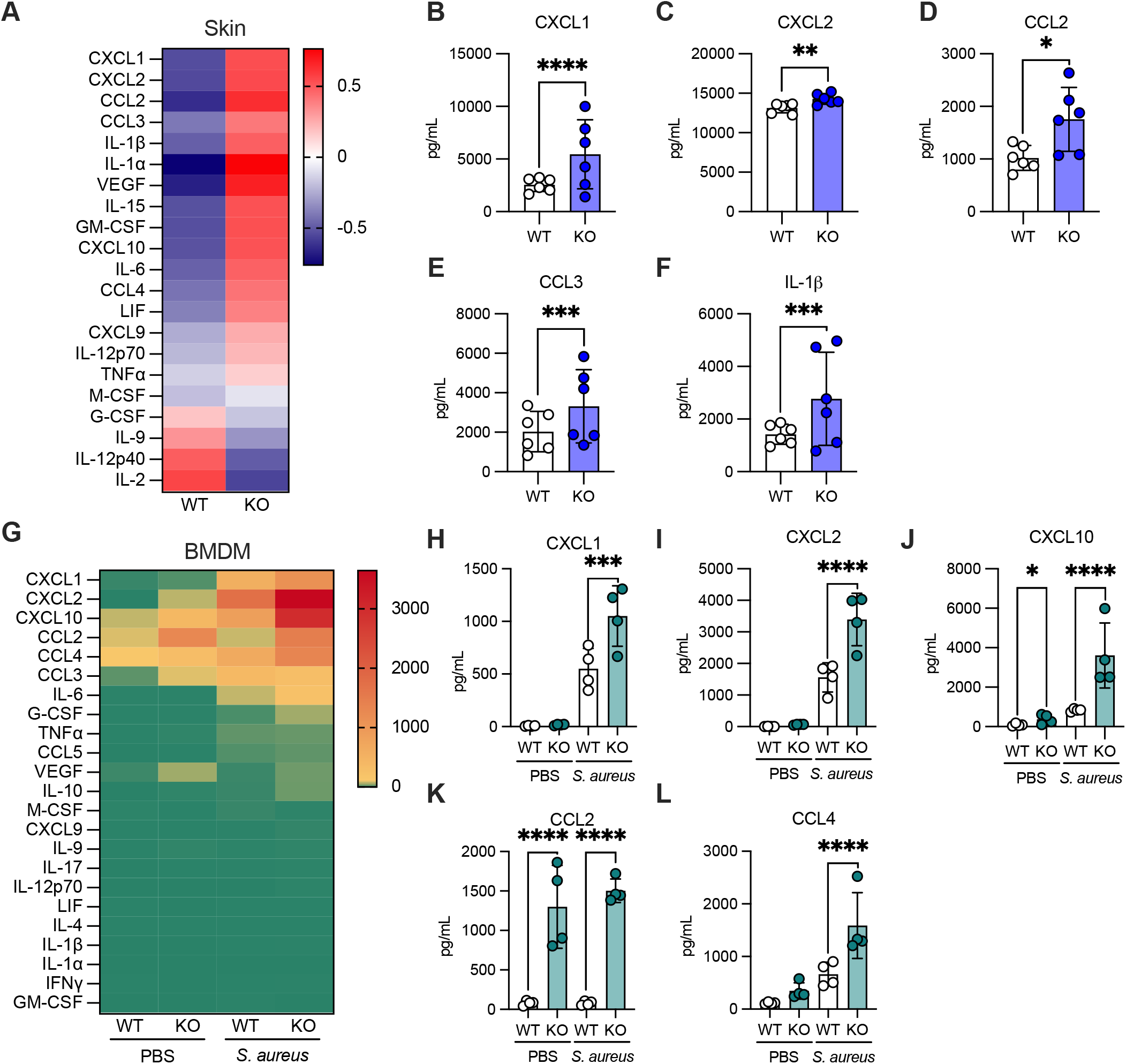
*Lair1* KO mice produce higher levels of pro-inflammatory cytokines. (**A**-**F**) Skin lesions were collected at 18 hours post-infection, homogenized, and cytokine production measured by multiplex cytokine array. (**A**) Heat map shows average WT and *Lair1* KO z-scored cytokine concentrations. (**B**-**F**) Bar graphs show all cytokines with significantly different expression between WT and *Lair1* KO. (**G**-**L**) Bone marrow-derived macrophages were generated in vitro and treated with PBS or *S. aureus* at a multiplicity of infection (MOI) of 10:1 for 18 hours. Cell supernatant was analyzed by multiplex cytokine array. (**G**) Heat map shows average WT and *Lair1* KO cytokine concentrations in PBS and *S. aureus* groups. (**H**-**L**) Bar graphs show all cytokines with significantly different expression between WT and *Lair1* KO. (n = 2 independent infections, 2-6 mice per group). Two-way ANOVA, Fisher’s LSD for multiple comparisons. *, p<0.05; **, p<0.01; ***, p<0.001; ****, p<0.0001

Because dermal macrophages express high levels of LAIR1 and are initial sensors of skin infection (14, 38), we hypothesized that macrophages were responsible for the pattern of increased cytokine production in skin infection at 18 hours post-infection. To test this hypothesis, we cultured bone marrow-derived macrophages (BMDMs) from WT and *Lair1* KO mice and measured cytokine production after PBS treatment or *S. aureus* infection using the same cytokine array method. We again observed a significantly increased overall pattern of cytokine expression in *Lair1* KO (**Fig. 3G**), with five significantly elevated chemokines, three of which were also increased in skin infection: CXCL1, CXCL2, and CCL2 (**Fig. 3H-L**). CXCL1 and CXCL2 were both increased by over 90% on average (**Fig. 3H-I**). Strikingly, CCL2 levels in *Lair1* KO were over ten-fold higher than WT in both PBS– and *S. aureus*-treated conditions (**Fig. 3K**). CXCL10 was also significantly higher at baseline in *Lair1* KO (**Fig. 3J**). Basal expression of several other chemokines and cytokines, including CCL3, CCL4, CXCL2, and VEGF, were non-significantly elevated at baseline in *Lair1* KO, suggesting that loss of LAIR1 releases inhibition of macrophage proinflammatory pathways at rest and may predispose cells to an excessive inflammatory response. Significant increases in pro-inflammatory cytokines that can recruit and activate multiple immune cell types, including monocytes and macrophages, were detected: CXCL10 was three-fold elevated (**Fig. 3J**) and CCL4 was elevated by over 130% (**Fig. 3L**) (39, 40). In summary, these data confirm our hypothesis that loss of LAIR1 inhibitory signaling in macrophages results in greater production of pro-inflammatory cytokines and support the notion that dermal macrophages are at least partially responsible for increased pro-inflammatory cytokine levels in *Lair1* KO skin abscess.

### Collagen and ECM remodeling pathways are elevated in Lair1 KO skin abscess and macrophages

Our results presented thus far demonstrate that *Lair1* KO macrophages produce high levels of proinflammatory mediators at baseline and in SSTI, leading us to hypothesize that the larger skin lesions and tissue damage in SSTI were due to excess inflammation. To begin to test this, we compared the transcriptomes of WT and *Lair1* KO *S. aureus* SSTI skin biopsies and WT and *Lair1* KO BMDMs treated with PBS or PAM3CSK4. PAM3CSK4 is a synthetic agonist of TLR2, which is stimulated in infection by bacterial lipoproteins, such as *S. aureus* lipotechoic acid, to induce NFκB pathways (41–43). We defined upregulated genes in RNAseq from *Lair1* KO BMDMs and *Lair1* KO *S. aureus* SSTI compared to WT and used GSEA to identify the top twenty pathways in these genes. Surprisingly, the top twenty Reactome pathways for both BMDMs and skin abscesses included four collagen-related extra-cellular matrix remodeling pathways: “collagen degradation”, “assembly of collagen fibrils and other multimeric structures” (denoted “collagen fibril assembly”), “collagen formation”, and “collagen biosynthesis and modifying enzymes” (denoted “collagen biosynthesis”) (**Fig. 4A-B**). Notably, there is substantial overlap in the genes in these pathways. Therefore, we combined them into a single set of 98 genes and plotted a heatmap of z-scored expression for BMDMs (**Fig. 4C**) and *S. aureus* SSTI (**Fig. 4D**) with genes grouped according to function: collagen, collagen synthesis, and ECM remodeling. In BMDMs and SSTI, *Lair1* KO exhibit a clear pattern of increased collagen-associated gene expression in response to PAM3CSK4 and *S. aureus*. Several collagen genes are also elevated in BMDMs at baseline (**Fig. 4C**), providing further evidence that loss of LAIR1 alters the macrophage phenotype prior to inflammatory insult.

**Figure 4.**
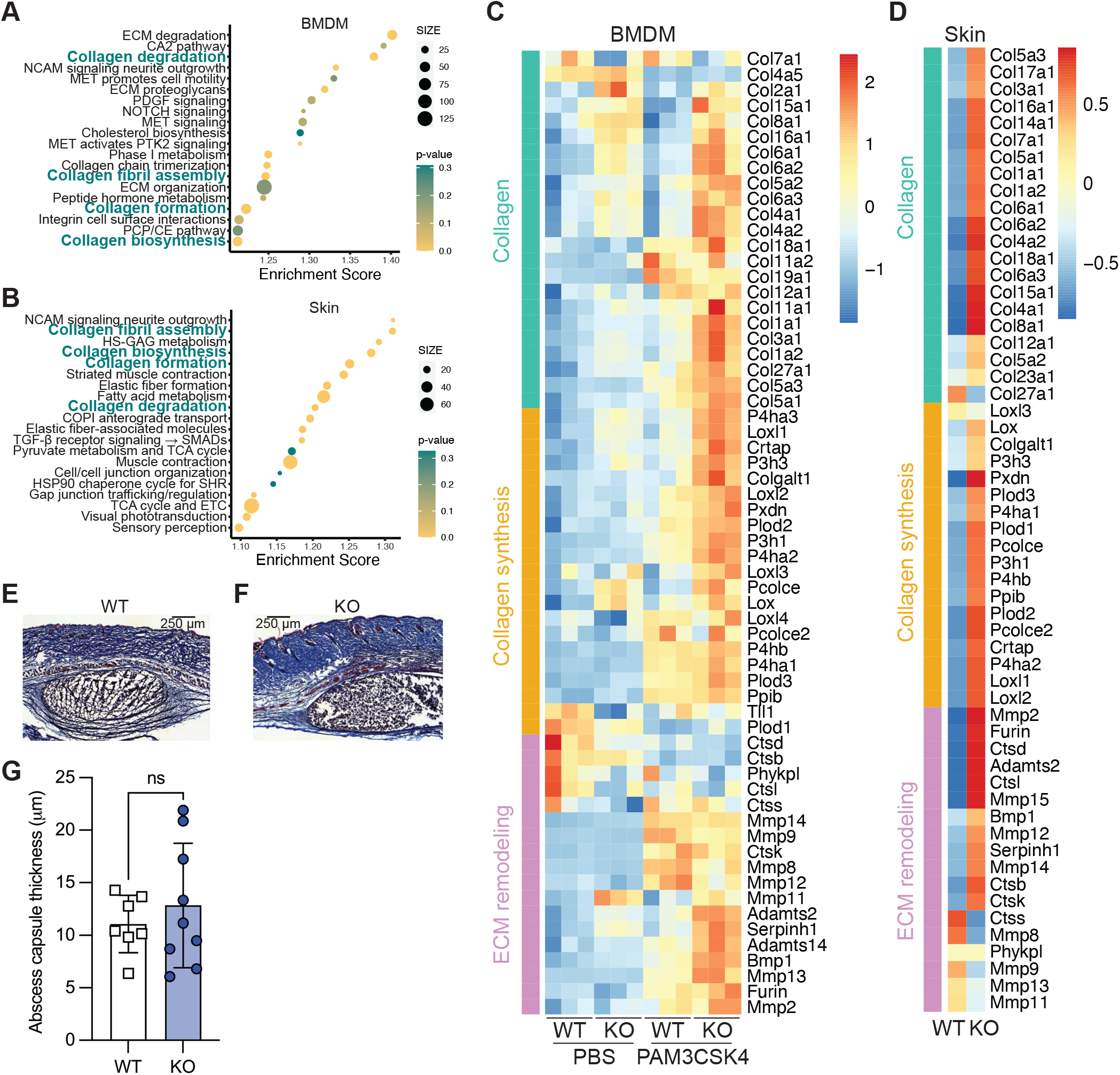
Collagen pathway gene expression is elevated in *Lair1* KO *S. aureus*-infected skin and macrophages. (**A**, **C**) Bone marrow-derived macrophages from WT and *Lair1* KO mice were treated with PBS or PAM3CSK4 for 6 hours, then subjected to RNA-sequencing. (**B**, **D**) Skin punch biopsies from WT and *Lair1* KO mouse *S. aureus* skin infection were taken at 2 dpi, then subjected to RNA-sequencing. GSEA was used to define the top 20 mouse Reactome pathways in genes upregulated in *Lair1* KO in BMDM treated with PAM3CSK4 (**A**) and skin infection (**B**); collagen-related pathways common to both analyses are bolded in teal. Genes from the four pathways, many of which overlapped, were classified into three general categories: collagen genes, collagen synthesis genes, and extracellular matrix (ECM) remodeling genes. Heat maps show z-scored expression for these genes for BMDM (**C**) and *S. aureus* skin infection (**D**). Example images from trichrome stained *S. aureus*-infected skin lesions at 1 dpi from WT (**E**) and *Lair1* KO (**F**) are shown, with abscess marked with an asterisk (dermonecrosis not yet developed at 1 dpi), as well as average capsule thickness at 3 points per lesion, measured in 4 WT and 5 KO skin lesions across multiple sections (**G**) (unpaired t-test, ns = non-significant).

Based on these findings, we hypothesized that differential collagen synthesis could lead to changes in abscess capsule formation in *Lair1* KO. To test this hypothesis, we compared frozen sections from WT (**Fig. 4E**) and *Lair1* KO (**Fig. 4F**) skin abscesses stained with Gomori’s trichrome, which stains collagen fibers. We measured average abscess capsule thickness over 3 points per lesion in 3 lesions from WT and *Lair1* KO and identified no difference (**Fig. 4G**). While we did not identify differences in wall thickness, the overall abscess area in *Lair1* KO was larger (**Fig. 1**), meaning total abscess wall circumference is correspondingly larger. The larger size would necessitate greater production of collagen and associated ECM remodeling enzymes to achieve, which is consistent with the RNAseq results. Collagens are also defense molecules and the increased expression of these pathways may be another facet of the enhanced inflammatory phenotype caused by loss of *Lair1*.

Because LAIR1 also binds complement proteins, which are involved in bacterial defense, we next investigated whether other LAIR1 ligands were also differentially regulated in *Lair1* KO abscess and BMDMs. We first evaluated the expression levels of collagen domain-containing complement proteins, including C1q subunits and lectins that are known LAIR1 ligands (10, 15). Of these, only *C1qa*, *C1qb*, and *C1qc* had detectable expression (averaged normalized counts > 100 for at least one group). In murine skin abscess, *C1qa* and *C1qc* levels were significantly higher in *Lair1* KO compared to WT (**Fig. S3A**). In BMDMs, *C1qa*, *C1qb*, and *C1qc* were also significantly higher in *Lair1* KO cells versus WT when stimulated with PAM3CSK4 (**Fig. S3B**). Together, these findings demonstrate that *Lair1* KO cells produce higher levels of collagens and collagen defense molecules when infected with *S. aureus* or stimulated by TLR2 ligands, consistent with their role in host immune response and with the notion that loss of LAIR1 causes a hyperinflammatory phenotype.

### Lair1 KO produce higher levels of reactive oxygen species but phagocytosis of S. aureus is equivalent

We hypothesized that *Lair1* KO neutrophils may also be hyperactivated in *S. aureus* infection because CXCL1 and CXCL2 were elevated in *S. aureus* infected skin. To test this hypothesis, we isolated neutrophils from WT and *Lair1* KO bone marrow and treated them with *S. aureus* and measured reactive oxygen species production. Compared to WT, *Lair1* KO neutrophils treated with *S. aureus* produced significantly more hydrogen peroxide (**Fig. 5A**), a major component of reactive oxygen species (ROS) critical for killing bacteria. Hydrogen peroxide can also cause extensive tissue damage (44, 45). We next assessed intracellular lysosome acidification because it is an important step in bacterial killing and SHP-1 plays a role in regulating this process(46, 47). We incubated neutrophils with *S. aureus* and Lys-NIR probe and measured fluorescence by flow cytometry. We did not detect any difference in lysosome acidification between WT and *Lair1* KO neutrophils (**Fig. 5B**). To determine whether there were differences in phagocytosis between WT and *Lair1* KO neutrophils, we incubated neutrophils with either *S. aureus* bioparticles (**Fig. 5C**) or live mCherry-expressing *S. aureus* (**Fig. 5D**), then measured *S. aureus* fluorescence in washed cells by flow cytometry. Neither measure was significantly different in WT and *Lair1* KO neutrophils.

**Figure 5.**
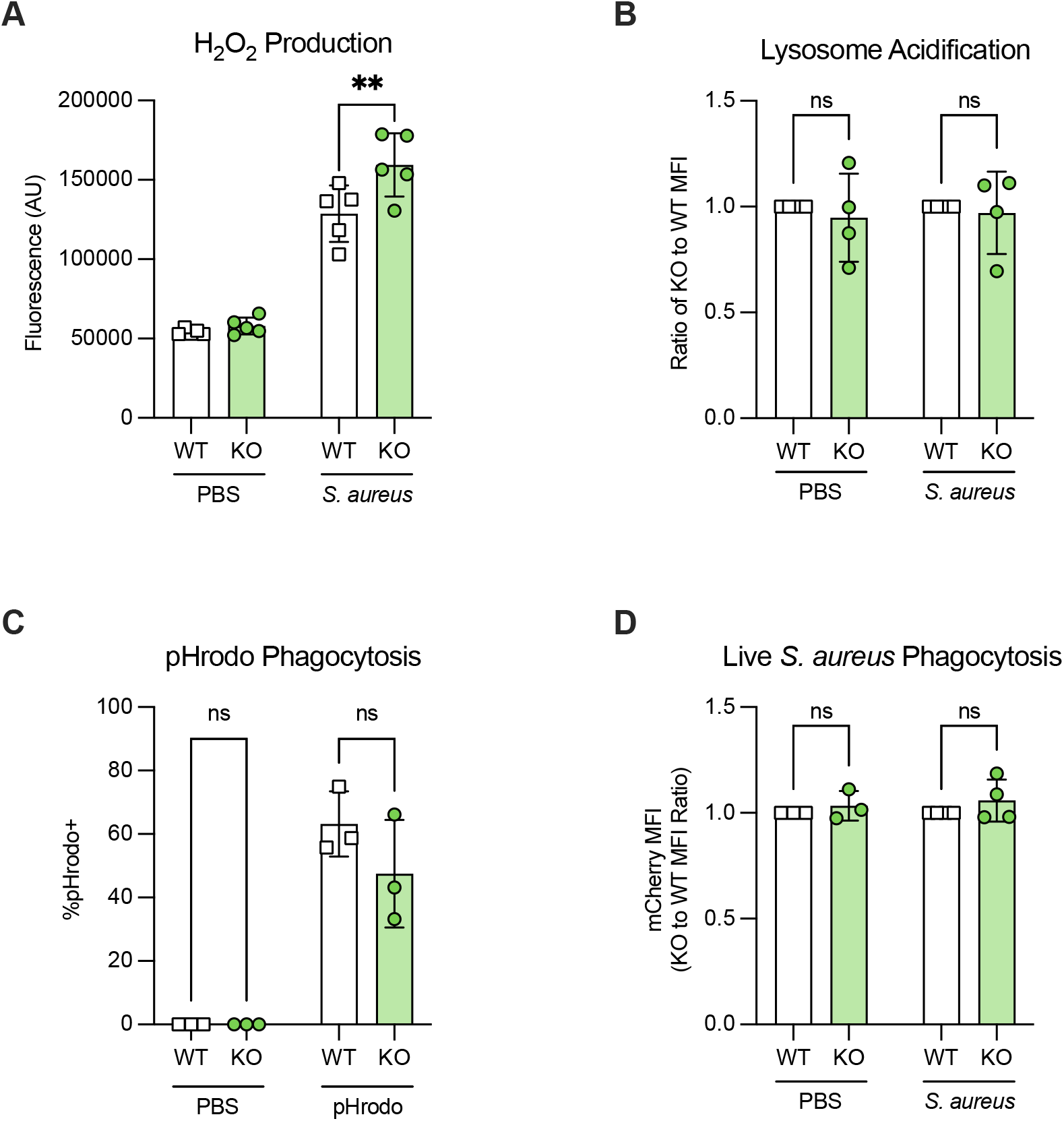
*Lair1* KO neutrophils produce increased reactive oxygen species but phagocytosis is equivalent to WT. WT and *Lair1* KO neutrophils were isolated from mouse bone marrow. (**A**) Amplex red fluorescence measured by plate reader indicates hydrogen peroxide (H_2_O_2_) production in neutrophils treated with PBS or *S. aureus* at MOI 50:1 for 30 minutes. (**B**) Ratio of WT to KO lysosome probe MFI measured by flow cytometry indicates lysosome acidification in neutrophils treated with PBS or *S. aureus* at MOI 100:1 for 30 minutes. (**C**) Neutrophils incubated for 30 minutes with PBS or *S. aureus* bioparticles (pHrodo) that fluoresce when acidified in the lysosome were subjected to flow cytometry. (**D**) Neutrophils incubated for 30 minutes with PBS or mCherry-expressing *S. aureus* at MOI 50:1 were subjected to flow cytometry. mCherry MFI is shown as a ratio of KO to WT. Significance (two-way ANOVA with Fisher’s LSD): ns, not significant; ** p < 0.01.

Monocyte-recruiting and –activating cytokines, particularly CCL2, were also elevated in *Lair1* KO (**Fig. 3**), therefore we also assessed lysosome acidification and phagocytosis in monocytes. Classical monocytes isolated from WT and *Lair1* KO bone marrow were assessed using the same methods as for neutrophils (**Fig. 5**). Like neutrophils, monocytes exhibited similar levels of phagocytosis and lysosome acidification in *Lair1* KO and WT (**Fig. S4A-B**). Together, these data demonstrate that *Lair1* KO neutrophils produce higher levels of ROS in SSTI, but neutrophil and monocyte phagocytosis and lysosome acidification are similar to WT. These data suggest that higher levels of ROS produce excess tissue damage in *Lair1* KO SSTI, yet do not enhance bacterial clearance.

### Cytokines elevated in Lair1 KO are also higher in advanced CTCL

To determine whether the cytokines that were elevated in *Lair1* KO were also increased in CTCL, we evaluated three publicly available RNA-seq datasets from CTCL lesional skin biopsies (7, 48, 49). We examined cytokine expression for all the significantly increased cytokines in *Lair1* KO BMDM and SSTI. Skin biopsies were collected from healthy donors and a range of CTCL stages, including: healthy donor, early stage CTCL, and late stage CTCL (**Fig. 6A**) (7); healthy donor, CTCL plaques, or CTCL tumors/patches (**Fig. 6B**) (48); or paired healthy and lesional skin from the same donors (**Fig. 6C**) (49). Notably, none of the biopsies contained tissue with active infection. CXCL1, CXCL10, CCL2 and CCL4 were significantly higher in at least 2 of 3 datasets, and CCL3 was significantly elevated in 3 of 3 datasets evaluated. CXCL2 and IL-1β were not elevated in any of the three datasets we evaluated, though other cytokines associated with CTCL pathogenesis were also elevated (**Fig. S5**). Overall, CTCL-involved skin exhibited a pattern of significantly increased expression of cytokines involved in myeloid cell recruitment and activation, though without infection-associated inflammatory signaling factors.

**Figure 6.**
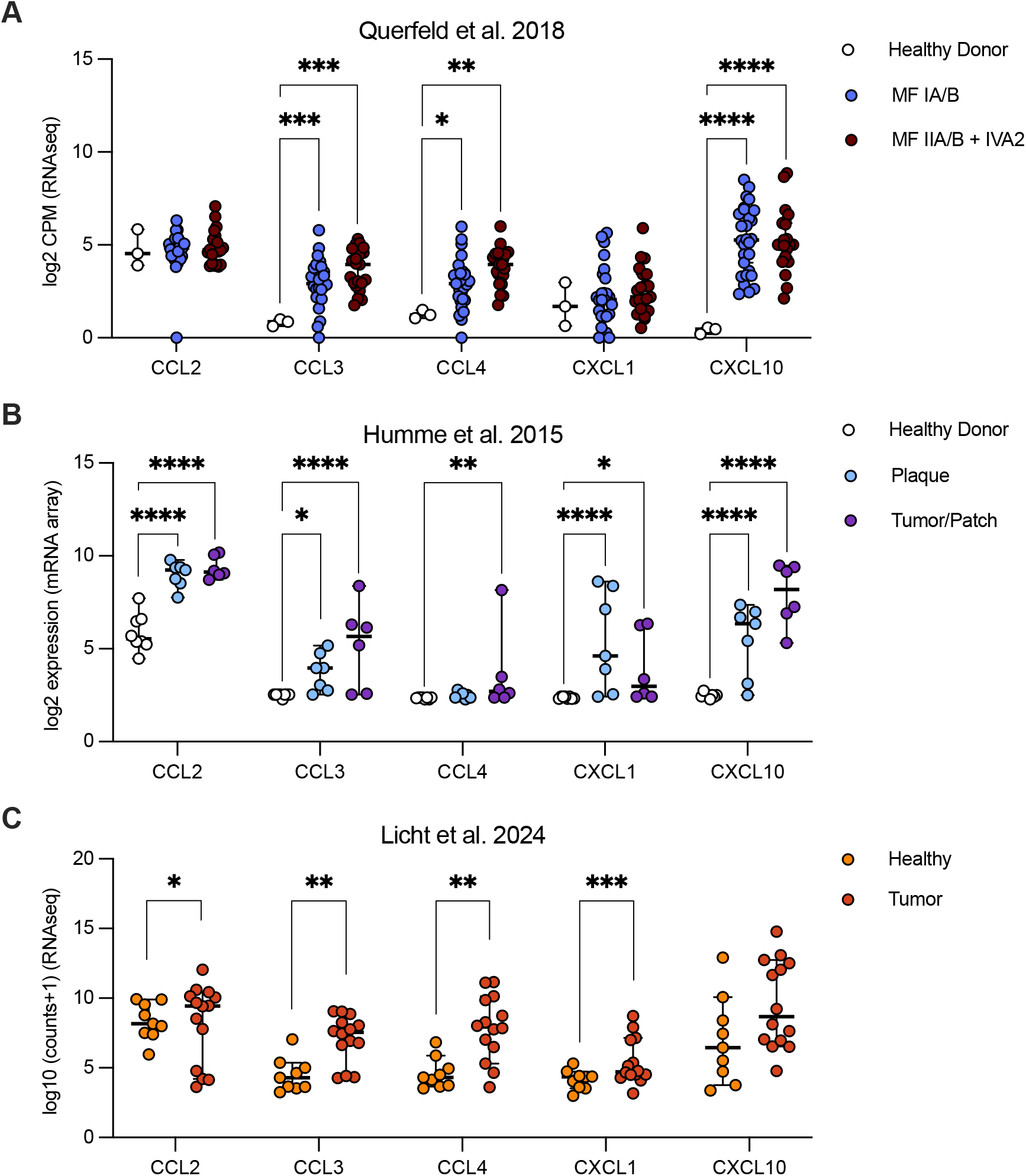
Cytokines elevated in *Lair1* KO are also higher in CTCL. Publicly available CTCL skin biopsy RNA-seq from Querfeld et al. 2018 (7) (**A**), Humme et al. 2013 (45) (**B**), and Licht et al. 2024 (46) (**C**) were analyzed using DEseq2. Normalized expression for CCL2, CCL3, CCL4, CXCL1, and CXCL10 is shown. Significance (DEseq adjusted p-value): *, p<0.05; **, p<0.01; ***, p<0.001; ****, p<0.0001

We next evaluated these three datasets for expression of the genes in collagen pathways that were elevated in *Lair1* KO *S. aureus*-infected skin and macrophages. Pathway analyses of upregulated genes identified significantly enriched gene sets in immune activation pathways such as CD28 co-stimulation, IL-3/IL-5/GM-CSF signaling, and IL-10 signaling, though no collagen/ECM remodeling pathways (**Fig. S6A-C**). However, we found that individual genes in these pathways were significantly increased: 5 ECM remodeling enzyme genes were elevated in 3 of 3 CTCL datasets (*CTSS*, *MMP1*, *MMP12*, *CTSB*, and *MMP9*) and 3 collagen genes were elevated in 2 of 3 CTCL datasets (*COL4A4*, *COL6A6*, *COLGALT2*) (**Fig. S6D-F**). These findings are not surprising, given that the biopsies did not involve active bacterial infection or abscess, but rather ongoing inflammation associated with the cutaneous lymphoma microenvironment (7, 48, 49). In summary, like *Lair1* KO *S. aureus* skin abscess and infected BMDMs, CTCL skin lesions exhibit an overall pattern of increased expression of inflammatory factors, including myeloid cell-recruiting and –activating cytokines and some collagen/ECM remodeling pathway genes.

## METHODS

### Sex as a biological variable

For human studies, both male and female subjects are included in approximately equal numbers. For animal studies, both male and female mice were used in equal proportions for *in vitro* experiments, including cytokine analysis from bone marrow-derived macrophage lysate, RNA sequencing of BMDMs, and experiments involving neutrophil and monocyte isolation. Male and female mice exhibit relevant differences in the stratum corneum – the surface epithelium of the skin. Visualization of epithelial lesions in these models is best performed in male mice as the stratum corneum is several cell layers thicker than in female mice. We confirmed that *S. aureus* skin infection in female mice exhibits a similar disease course as in male mice, including differences in *Lair1* KO vs WT. Tissue-based analysis of the clinical correlates of disease are less informative in female mice, due to the very thin stratum corneum. Therefore, to minimize the number of animals needed to obtain consistency in the *S. aureus* skin infection models, only male mice were used, which is in agreement with previous reports on this model (50).

### Mice

Wild type C57BL/6J and *Lair1*^-/-^ (B6.Cg-Lair1tm1.1Jco/J, JAX Strain #:032788) mice(28) were purchased from The Jackson Laboratory (ME, USA) and housed in a pathogen-free environment in the WUSM animal facility. Breeders were fed high fat chow, and all other mice were fed standard rodent chow (5058; Purina, St. Louis, MO, USA). Inhaled isoflurane was used for all anesthesia.

### Bacteria

*Staphylococcus aureus* (*S. aureus*) MRSA strain USA300/LAC was used for all infections. Colonies from tryptic soy agar plates were grown overnight at 37° C in tryptic soy broth, then subcultured at a 1:10 dilution and grown until 0.5-0.7 OD_600_. Culture was diluted with tryptic soy broth to OD_600_ 0.48, then bacteria were pelleted, washed in PBS, and diluted to the desired inoculum using the estimated dilution factor 2×10^8^ bacteria/mL. Cultures were serially diluted and then plated onto tryptic soy agar to confirm desired inoculum.

### Skin Infections

On day 0, two-to three-month-old male mice were anesthetized, then flanks were shaved and treated with hair removal cream for 2-3 minutes. After 24 hours, flanks were injected subcutaneously with expected dose of 1×10^7^ CFU *S. aureus* (actual dose ranged from 1-2×10^7^ CFU). Abscess and dermonecrosis length and width were measured over 14 days using digital calipers, then area of abscess and dermonecrosis was calculated using the formula A = ([pi/2] x l x w). Dedicated mice were sacrificed at 24 or 48 hours after infection for 8mm skin punch biopsy. For histology, skin punch biopsies were fixed in 10% neutral-buffered formalin for 24 hours at 4°C for paraffin-embedding and H&E staining (Washington University) or fixed for 2 hours in 10% neutral-buffered formalin at room temperature before embedding in O.C.T. Compound (Fisher, 4585) prior to trichrome and H&E staining (WUSM Pulmonary and Critical Care Morphology Core). For CFU analysis or cytokine production analysis, skin punch biopsies were homogenized in PBS and either serially diluted and plated onto tryptic soy agar (31) or stored at –80°C until analysis, respectively. For RNA-sequencing, skin punch biopsies were homogenized in TRI reagent (Sigma, 93289) and stored at –80°C until RNA isolation. Results from an entire 8mm skin punch biopsy are reported for each analysis.

### Preparation of primary cells from mouse bone marrow

Bone marrow from age– and sex-matched WT and Lair1^-/-^ mice was used for all primary cells. Bone marrow-derived macrophages (BMDMs) were produced by treating bone marrow cells with the equivalent of 10 ng/mL M-CSF from CMG14-12 cells (Veis lab, Washington University in St. Louis) for 6 days. Primary monocytes and neutrophils were isolated from mouse bone marrow using negative isolation kits (Miltenyi Biotec, 130-100-629 and 130-097-658). Cells were cultured in RPMI 1640 media (monocytes, neutrophils) or a-MEM media (BMDMs) supplemented with 10% FBS, 100 U/mL penicillin (Corning), 100 mg/mL streptomycin (Corning), and 50 mM ®-mercaptoethanol at 37° C in 5% CO_2_.

### In vitro assays

Amplex Red Hydrogen Peroxide Assay Kit (Invitrogen, A22188) was used according to manufacturer’s instructions to measure reactive oxygen species production. To measure phagocytosis and lysosome acidification, neutrophils and monocytes were treated with either pHrodo Green *S. aureus* BioParticles (Invitrogen, P35382) or mCherry-expressing *S. aureus* (gift from the Veis lab, Washington University) (29) at multiplicity of infection 50:1 with or without lysosome probe NIR (Biolegend, 421916) at a concentration of 1:500. After incubation at 37° C in 5% CO_2_ for 30 minutes, cells were immediately transferred to ice and pHrodo, mCherry, or lysosome probe fluorescence analyzed by flow cytometry.

### Cytokine analysis

Multiplex cytokine array (32-plex discovery panel, Eve technologies) was conducted for skin homogenate and BMDM supernatant.

### Patient sample collection

De-identified peripheral blood was obtained from patients seen at the Washington University School of Medicine Cutaneous Lymphoma Clinic under IRB-approved protocols with patients providing informed consent.

### Peripheral blood mononuclear cell isolation and scRNA-sequencing

PBMCs were subjected to enrichment and/or depletion using antibody cocktails (Miltenyi Biotec) to enable purification of the desired cells. Monocytes and neutrophils were depleted by incubation of peripheral blood samples with RosetteSep Human Monocyte (CD36) Depletion Cocktail for 15 minutes at room temperature, layered onto a Histopaque-1077 gradient, and centrifuged at 400g for 30 minutes with no brake. The interphase was collected and washed with 10 mL of sort buffer (PBS, 1% FBS, 2 mM EDTA), followed by red blood cell lysis (155 mM NH_4_Cl, 10 mM KHCO_3_, 0.1 mM EDTA) for 10 minutes. For each sample, 10,000 to 20,000 viable cells were submitted for processing using the 10x Genomics Chromium Controller and the Chromium Single Cell 5′ Library & Gel Bead Kit v2 (PN-1000006), Chromium Single Cell A Chip Kit (PN-1000152), Chromium Single Cell V(D)J Enrichment Kit, Human, Tcell (96rxns)(PN-1000005), and Chromium Single Index Kit T (PN-1000213) following the manufacturer’s protocols. Normalized libraries were sequenced on a NovaSeq6000 S4 Flow Cell using the XP workflow and a 151×10×10×151 sequencing recipe according to manufacturer protocol. A median sequencing depth of 50,000 reads/cell was targeted for each sample.

### scRNA-sequencing data processing and analysis

Alignment and gene counting were performed using the Cell Ranger pipeline (10x Genomics, v3.0, Pleasanton, CA). Genes found in fewer than 15 cells in a given sample were removed. For each patient, gene counts and cells were pooled into a single Seurat (v3.1.4) object (51), and cells containing fewer than 200 or more than 3000-3750 expressed genes, more than 8-10% mitochondrial reads, fewer than 300 or more than 10000-20000 UMIs, or classification as a doublet by the R package scDblFinder with parameters dims = 30, clust.method = “fast_greedy” were removed. Normalization and regression of technical variation due to mitochondrial read percentage and read depth was performed with the SCTransform function with variables.features.n = 4000. Integration to account for experimental variability due to differences in ficoll or buffy coat preparation and batch effects was performed using the Seurat wrapper around the fastMNN function from the batchelor R package (v1.4.0) with n.features = 3000. Gene expression was normalized and the top 1500 variable using the “VST” method were calculated. Data was integrated using the harmony (v1.0.0) R package (52) using both patient and batch information to correct for batch effect with up to 10 iterations. The UMAP and neighbors were calculated with Seurat, using 20 dimensions of the harmony calculations. Cell annotation was performed using the singler (v1.4.1) R package (53) using the highest spearman rho of purified immune populations in the Human Primary Cell Atlas (34). Cell type designations with less than 50 cells in the entire cohort were reduced to “other”. Automated annotations were checked manually using canonical marker genes. All single cell visualizations were performed with Seurat and ggplot2 R package (v3.5.1) (54).

### RNA isolation and sequencing

RNA was isolated from homogenized mouse skin punch biopsies using a Direct-zol RNA MiniPrep Plus kit (Zymo research, R2072). Bone marrow-derived macrophages were plated at 2×10^5^ cells/mL and treated with PBS or 500 ng/mL PAM3CSK4 (Tocris Bioscience, 4633) for 6 hours. RNA was isolated using an RNeasy Plus Mini kit (Qiagen, 74134). RNA was sequenced by the Genome Technology Access Center at the McDonnell Genome Institute at Washington University School of Medicine on an Illumina NovaSeq-6000 to generate 150 bp paired end reads. RNA-seq reads were then aligned to the Ensembl release 101 primary assembly with STAR (v 2.7.9a1). Gene counts were derived from the number of uniquely aligned unambiguous reads by Subread:featureCount (v 2.0.32). Isoform expression of known Ensembl transcripts were quantified with Salmon (v 1.5.23).

### RNA-seq analysis

R (v 4.1.3) was used for all analyses. DEseq2 (v 1.4.1) (55) was used to normalize gene counts and evaluate significance of individual genes for our own RNAseq and publicly obtained datasets. Genes with counts above expression cutoffs (averaged normalized counts > 100 for at least one group), significance < 0.5, and absolute log_2_ fold change > 0.1 were analyzed by GSEA (v 4.3.2) (56, 57) desktop software using the MSigDB mouse or human reactome (v.2023.2) gene set for pathway analysis. Heatmaps were generated using pheatmap (v 1.0.12) R package.

### Statistical Analyses

GraphPad Prism 9 (Version 9.4.1) was used for all statistical tests other than RNA-seq.

### Study Approval

De-identified peripheral blood draws were obtained from patients seen at the Washington University School of Medicine Cutaneous Lymphoma Clinic under IRB-approved protocols with patients providing informed consent. All animal experimental protocols were approved by Institutional Animal Care and Use Committee of Washington University in St. Louis. All animal experiments were performed in accordance with National Institutes of Health Guide for the Care and Use of Laboratory Animals.

### Data Availability

Sequencing data was deposited in GEO under accession numbers GSE269177 and GSE269178. External datasets used include GEO datasets GSE59307, GSE113113, and GSE221148.

## DISCUSSION

Our studies presented here reveal new roles for LAIR1 inhibitory signaling in CTCL and bacterial infection. In CTCL, a cutaneous cancer with inflammatory damage to skin, loss of LAIR1 signaling due to excessive LAIR2 is associated with high levels of pro-inflammatory cytokine expression and advanced disease. In *S. aureus* infection, which is common in CTCL, we discovered that LAIR1 regulates the host immune response, and that loss of LAIR1 causes greater inflammation and tissue damage.

We previously identified high levels of LAIR2, which is a decoy receptor for LAIR1 and blocks its activity, in CTCL patients with progressive disease (8). CTCL progression is marked by skin inflammation and barrier breakdown, which culminate in frequent *S. aureus* skin and soft tissue infections. LAIR1 had been previously shown to dampen activation in diverse immune cell subsets (9, 12, 18, 58). Therefore, we hypothesized that loss of LAIR1 inhibitory signaling may promote increased inflammation in CTCL skin lesions. Inflamed skin has long been recognized as a risk factor for *S. aureus* infection, particularly in CTCL (59, 60). Thus, we decided to test LAIR1 loss in a well-established mouse model of *S. aureus* SSTI (37, 50). We observed a striking phenotype in *Lair1* KO mice: they suffered larger skin lesions with more tissue damage and higher levels of pro-inflammatory cytokines in *S. aureus* SSTI.

Although LAIR2 is produced primarily by NK and T cells, including CTCL tumor cells, it is secreted and therefore can block LAIR1 signaling on other cells (8, 10, 61). Innate immune cells, including monocytes and macrophages, express high levels of LAIR1 (13), and thus would be most affected by LAIR2 blockade. Related to this, innate immune cells are the first responders in acute bacterial infection. Therefore, we predicted that loss of inhibitory signaling in these cells would result in an early and excessive inflammatory response to acute bacterial infection. Consistent with our prediction, our studies here show that *Lair1* KO exhibit increased pro-inflammatory cytokine levels and larger areas of abscess and dermonecrosis just 1-2 days post *S. aureus* infection. While other studies showed an impact of LAIR1 on viral and malarial infection (24, 25, 27), our studies are the first to our knowledge to demonstrate that LAIR1 plays a role in modulating inflammatory response to bacterial infection.

Pro-inflammatory cytokine production in *S. aureus*-infected *Lair1* KO BMDMs mirrored that of infected skin, suggesting that skin macrophages may be responsible, at least at early infection time points. This notion is further supported by the fact that most are myeloid chemokines, including CXCL1 and CXCL2, which serve to recruit neutrophils from the peripheral blood to the site of infection, resulting in abscess formation (62). Indeed, *Lair1* KO had increased abscess size, likely due to increased neutrophil recruitment and influx. These results are consistent with a study in *Lair1* KO lung that showed increased neutrophil recruitment upon oropharyngeal administration of CXCL1 and RSV infection compared to WT, although RSV infection did not cause increased cytokine production in *Lair1* KO in this study (63). Our findings thus demonstrate a novel role for LAIR1 in regulating cytokine production and neutrophil recruitment in bacterial infection.

Given these results, we next sought to determine if loss of LAIR1 alters relevant functions of neutrophils or monocytes. Reactive oxygen species (ROS) production was higher in *Lair1* KO neutrophils compared to WT, though phagocytic ability in neutrophils and monocytes was equivalent to WT. ROS production causes bacterial as well as host cell death, and if unchecked, leads to significant tissue injury (64, 65). The larger areas of dermonecrosis in *Lair1* KO may be due, at least in part, to increased ROS production.

Another novel finding of this study is the increased expression of complement, collagens, and ECM remodeling genes in *Lair1* KO SSTI and BMDMs. Complement proteins are important in host defense against microbial pathogens (66). Collagens and enzymes involved in production and degradation of the ECM are essential for abscess formation (62). Complement component C1q and collagen proteins are ligands of LAIR1. These findings raise the intriguing possibility of a novel role for LAIR1 in the setting of infection, in which collagen breakdown and complement consumption decrease the pool of LAIR1 ligands, which reduces LAIR1 inhibitory signaling, which in turn increases production of C1q and collagen via a feed-forward mechanism.

Our studies revealed several similarities in CTCL biopsy tissue and *Lair1* KO *S. aureus* infection. Both exhibit increases in macrophage– and monocyte-activating cytokines, including CXCL10, which has previously been linked to early CTCL pathogenesis (67–69). Neutrophil infiltration in CTCL is rare (70, 71) and, accordingly, we did not find similar increases in neutrophil activating cytokines in CTCL. Notably, the CTCL skin biopsies evaluated in this study did not harbor active *S. aureus* infection, which likely explains this lack of overlap. Overall, our results support that inhibition of LAIR1 signaling via LAIR2 overexpression causes macrophage-initiated inflammatory tissue damage and *S. aureus* susceptibility in CTCL.

Particularly intriguing are our findings that some pro-inflammatory cytokines and collagen pathway genes have higher basal expression levels in *Lair1* KO macrophages. These results suggest that loss of LAIR1 may cause macrophages to be “pre-activated”, i.e., exhibit some characteristics of activation prior to immune stimulus, which could be indicative of altered regulation of immune response at the epigenetic level. These observations require further study to elucidate the underlying mechanisms.

Together, our studies reveal LAIR1 as a critical modulator of pathogenic inflammation that is important for immune response to *S. aureus* infection and protection from CTCL immunopathology. We additionally demonstrate that LAIR1 loss increases the expression of cytokine signaling pathways, which has broader implications for other inflammatory diseases and malignancies in which LAIR1 signaling has been implicated (21, 72–75). Finally, our findings raise exciting new questions regarding the impact of LAIR1, and other inhibitory immune receptors, on the regulation of genes involved in inflammation. Future studies are needed to investigate these compelling new ideas.

## Acknowledgements

We thank the Alvin J. Siteman Cancer Center at Washington University School of Medicine and Barnes-Jewish Hospital in St. Louis, MO. and the Institute of Clinical and Translational Sciences (ICTS) at Washington University in St. Louis, for the use of the Genome Technology Access Center, which provided library preparation, sequencing, and data analysis services (detailed in Methods). The Siteman Cancer Center is supported in part by an NCI Cancer Center Support Grant #P30 CA091842 and the ICTS is funded by the National Institutes of Health’s NCATS Clinical and Translational Science Award (CTSA) program grant #UL1 TR002345.

We acknowledge the Genome Technology Access Center at the McDonnell Genome Institute at Washington University School of Medicine for help with genomic analysis. The Center is partially supported by NCI Cancer Center Support Grant #P30 CA91842 to the Siteman Cancer Center from the National Center for Research Resources (NCRR), a component of the National Institutes of Health (NIH), and NIH Roadmap for Medical Research. We also acknowledge the Research Infrastructure Services (RIS) group at Washington University in St. Louis for providing computational resources and services needed to generate the research results delivered within this paper. URL: https://ris.wustl.edu.

We thank Ezri Perrin for assistance with editing.

## Author Contributions

Conception and design: H.K.D., D.J.V., J.B.W., J.E.P. Development of methodology: H.K.D., K.T., E.C.M., J.E.P. Acquisition of data: H.K.D., K.T., E.C.M., L.T., A.C.M., N.M.S., C.C.Q., M.K.S., J.E.P. Analysis, interpretation of data, generation of figures: H.K.D., K.T., J.E.P. Funding acquisition, writing, review, and/or revision of the manuscript: H.K.D., E.C.M, N.M.S., J.E.P. Study supervision: J.E.P. All authors reviewed and approved the manuscript.

## Funding

This work was supported by American Heart Association Predoctoral Fellowship grant #P23-01044 to H.K.D.; Washington University BioSURF funding to E.C.M.; Siteman Cancer Center Foundation, Barnes-Jewish Hospital Foundation Steinbeck Designated Fund, and McDonnell Genome Institute and Illumina research support to N.M.S. and J.E.P.

## Conflict of Interest Statement

**The authors declare no conflict of interest.**

**Supplementary Figure 1.**
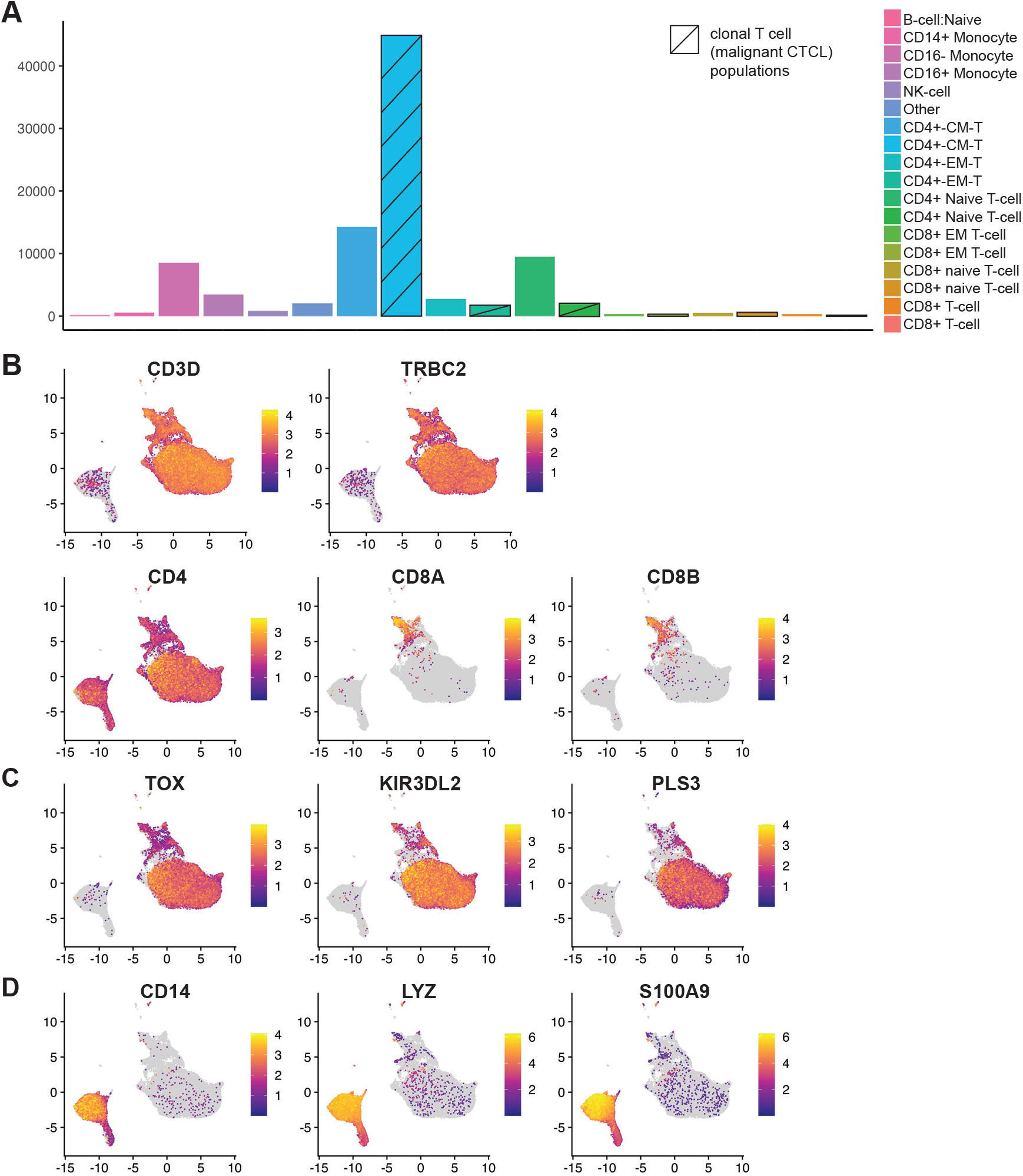
scRNAseq cell type distribution and canonical marker genes demonstrates separation of benign and malignant T cells. (**A**) Distribution of cell types based on HPCA cell annotation analysis for 92,496 mononuclear cells from 16 PBMC samples from 6 CTCL patients. (**B**-**D**) UMAP projection all cells from (**A**) shows expression of selected canonical T-cell (**B**), CTCL (**C**), and myeloid (**D**) genes.

**Supplementary Figure 2.**
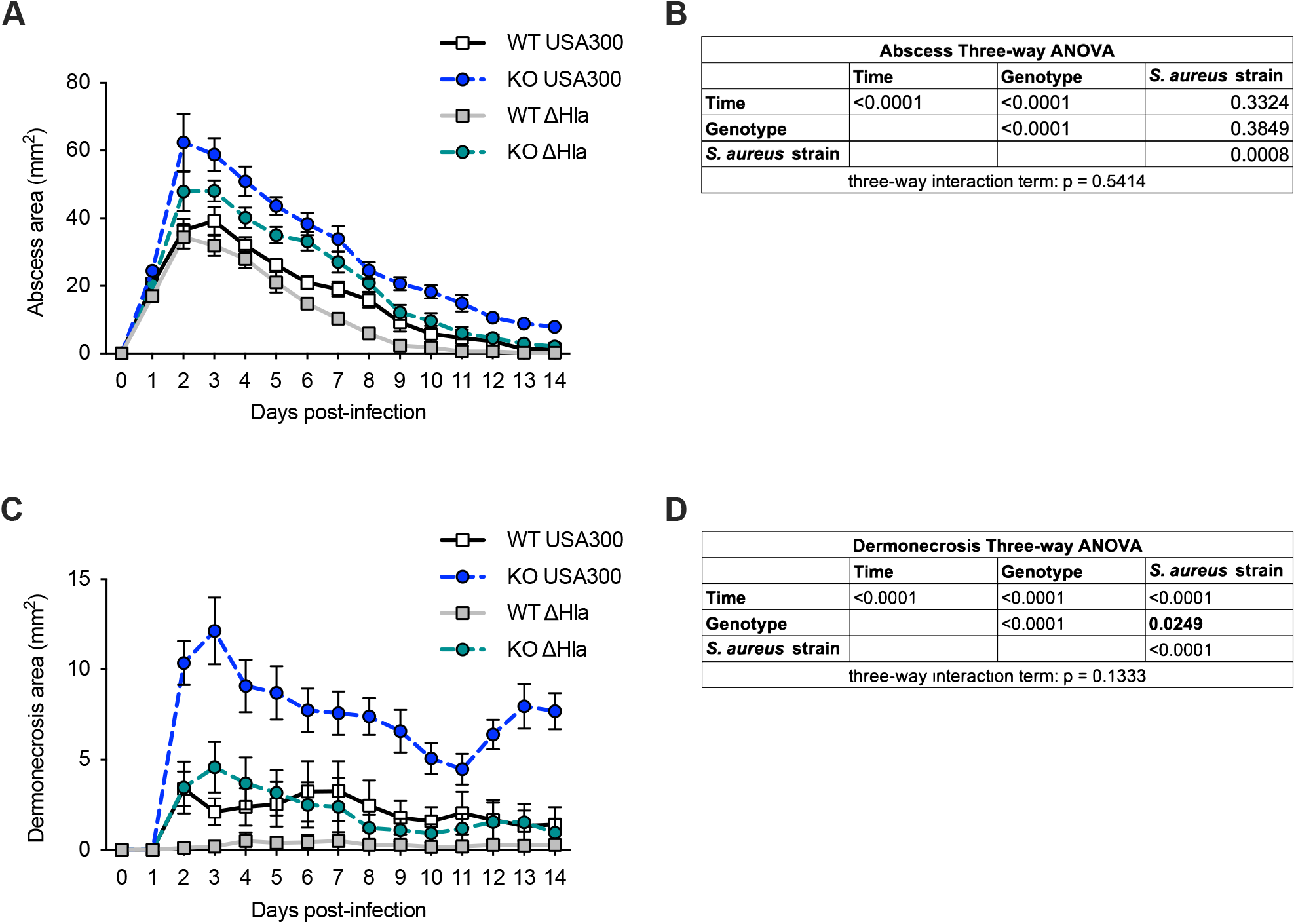
Increase in abscess and dermonecrosis size in *S. aureus* skin infection in *Lair1* KO is maintained in the absence of Hla. WT and *Lair1* KO mice were infected on opposite flanks with USA300/LAC and isogenic Hla knock-out strain (Δ Hla), then lesion size measured over 14 days. Abscess area over 14 days (**A**) and corresponding hypothesis testing (**B**) shown above; dermonecrosis area over 14 days (**C**) and corresponding hypothesis testing (**D**). Three-way ANOVA tables show p-values for single variable effects shown on the diagonal, two-way interaction effects shown between respective variables, and three-way interaction effect specified below (**B**, **D**), with significant interaction terms involving genotype bolded.

**Supplementary Figure 3.**
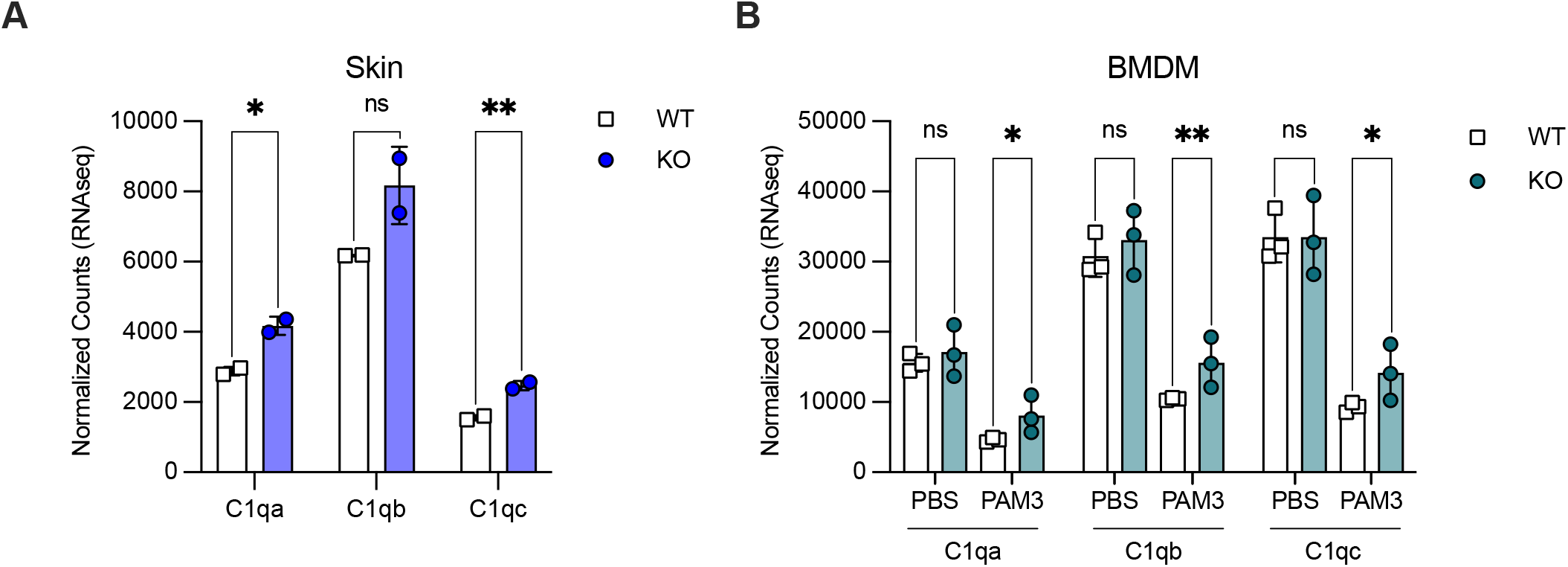
Higher complement C1q in *Lair1* KO *S. aureus*-infected skin and BMDM. Normalized gene counts for C1qa, C1qb, and C1qc in WT and *Lair1* KO *S. aureus*-infected skin (**A**) and BMDMs stimulated with PBS or PAM3CSK4 (PAM3) (**B**). Significance (DESeq2 adjusted p-value): ns, not significant; *, p < 0.05; **, p<0.01.

**Supplementary Figure 4.**
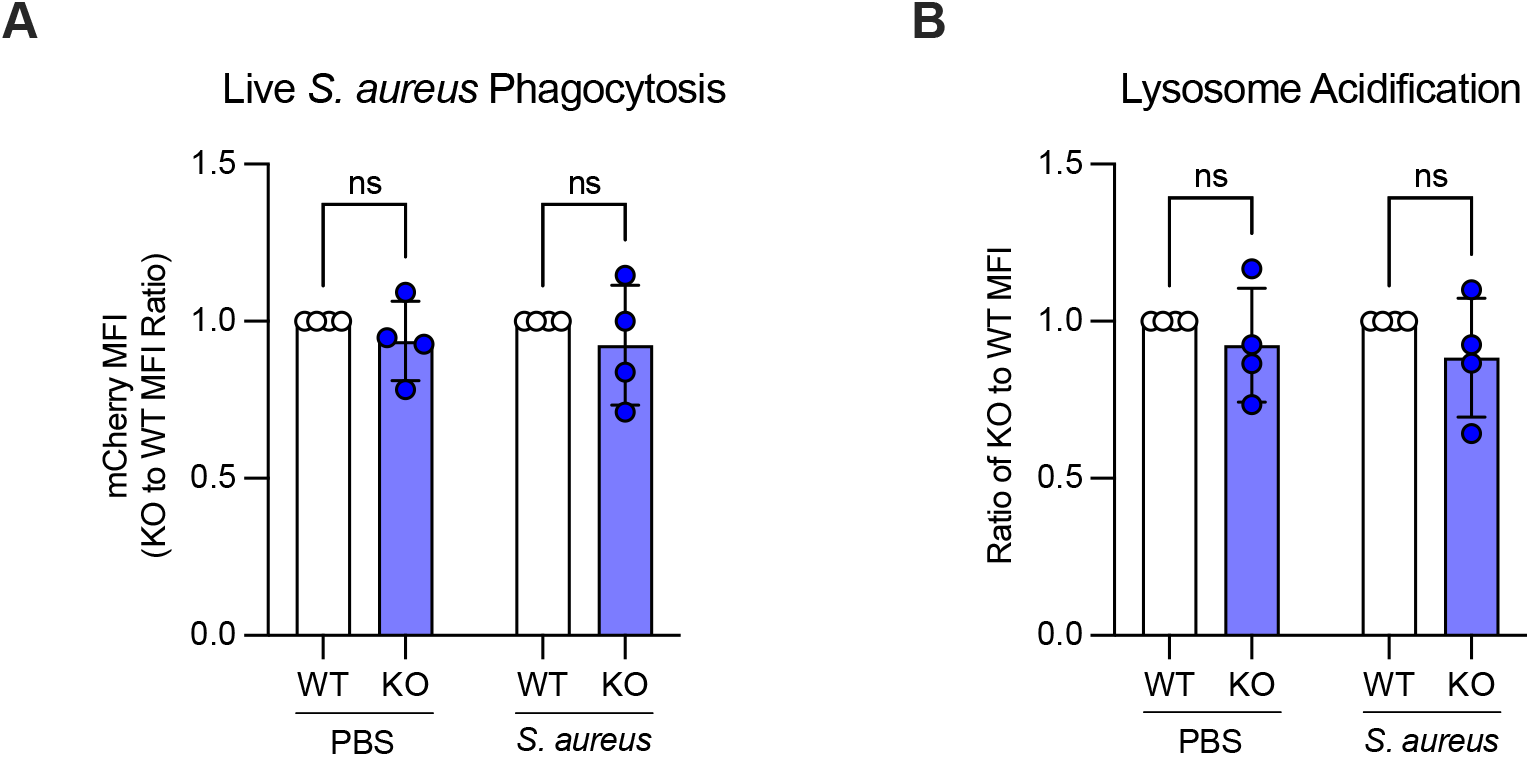
*Lair1* KO and WT monocytes exhibit equivalent phagocytosis of *S. aureus*. Classical monocytes were isolated from WT and *Lair1* KO bone marrow. (**A**) Monocytes were incubated for 30 minutes with PBS or mCherry-expressing *S. aureus* at MOI 50:1 and subjected to flow cytometry. mCherry MFI is shown as a ratio of KO to WT fluorescence. (**B**) Ratio of WT to KO lysosome probe MFI measured by flow cytometry indicates lysosome acidification in monocytes treated with PBS or *S. aureus* at MOI 100:1 for 30 minutes. Significance (two-way ANOVA with Fisher’s LSD): ns, not significant

**Supplementary Figure 5.**
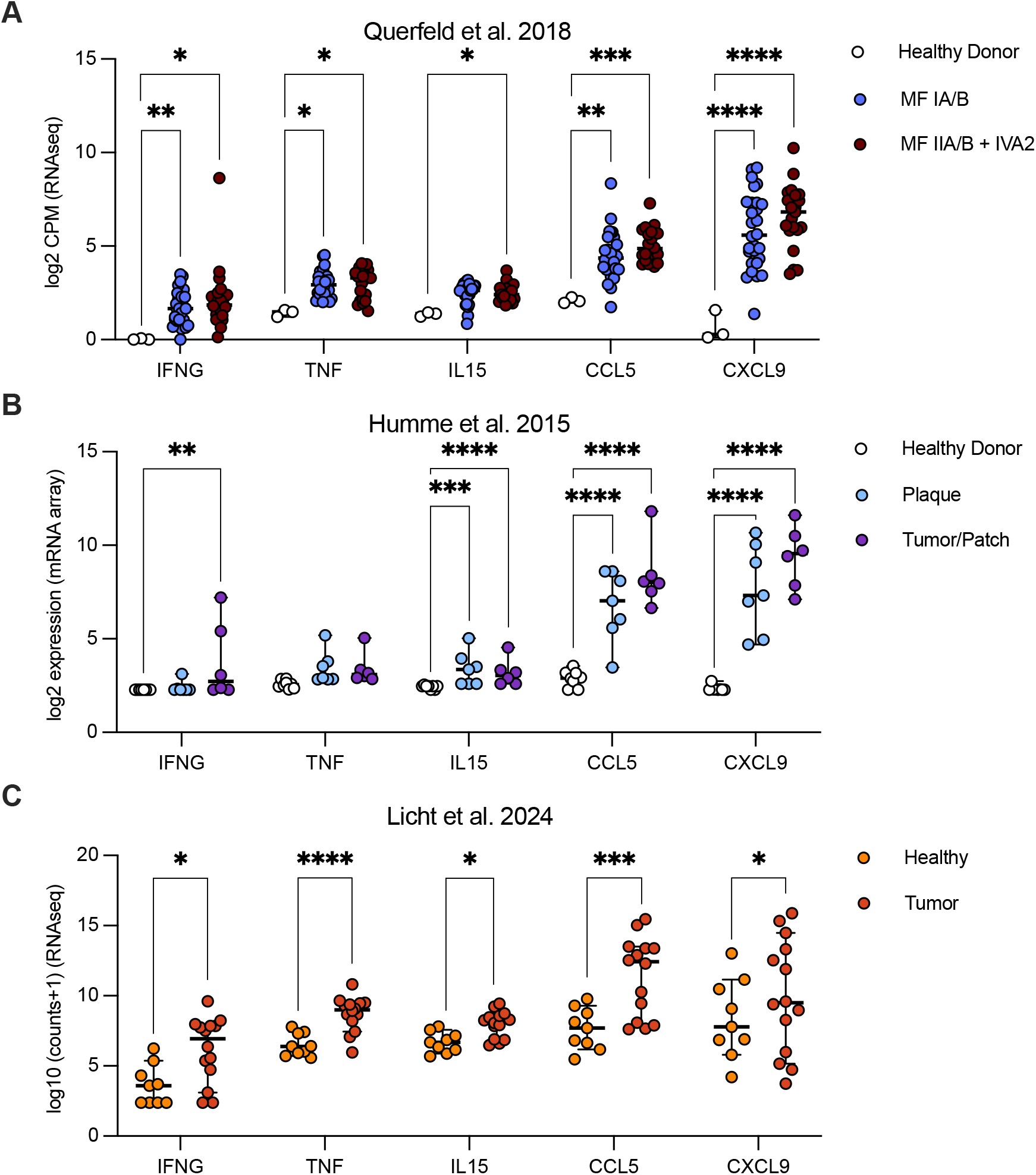
Some cytokines elevated in CTCL are not elevated in *Lair1* KO. Publicly available CTCL skin biopsy RNA-seq from Querfeld et al. 2018 (7) (**A**), Humme et al. 2013 (45) (**B**), and Licht et al. 2024 (46) (**C**) were analyzed using DEseq2. Normalized expression for IFNG, TNF, IL15, CCL5, and CXCL9 is shown. Significance (DEseq adjusted p-value): *, p<0.05; **, p<0.01; ***, p<0.001; ****, p<0.0001

**Supplementary Figure 6.**
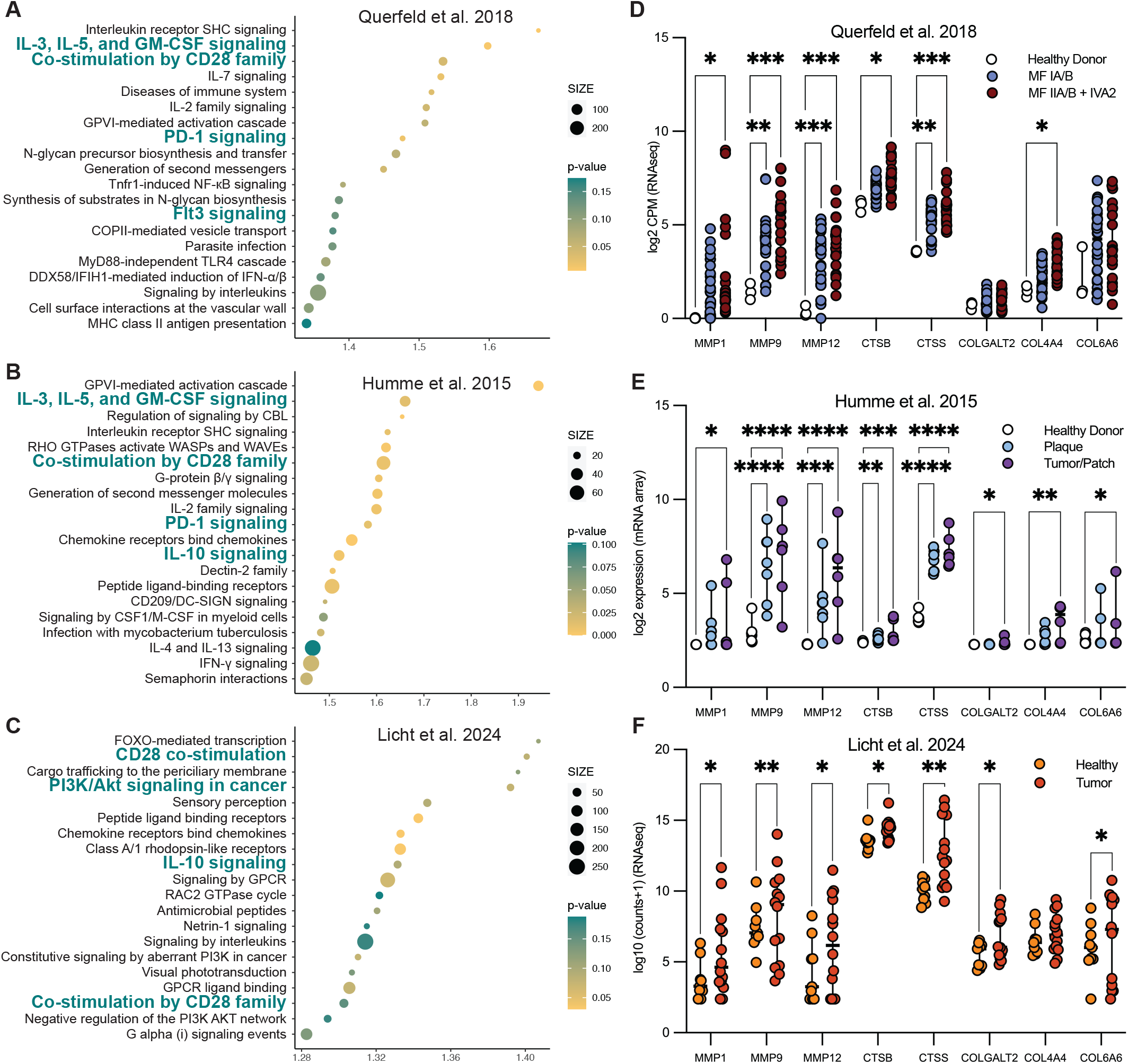
Several collagen and ECM remodeling genes are elevated in CTCL. Publicly available CTCL skin biopsy RNA-seq from Querfeld et al. 2018 (7) (**A**, **D**), Humme et al. 2013 (45) (**B**, **E**), and Licht et al. 2024 (46) (**C**, **F**) were analyzed using DEseq2 and GSEA. Top twenty pathways elevated in CTCL by GSEA human reactome analysis (**A**-**C**) and normalized expression for collagen pathway genes significantly increased in at least two datasets, including MMP1, MMP9, MMP12, CTSB, CTSS, COLGALT2, COL4A4, and COL6A6, are shown (**D**-**F**). Significance (DEseq adjusted p-value): *, p<0.05; **, p<0.01; ***, p<0.001; ****, p<0.0001

## Notes

https://www.ncbi.nlm.nih.gov/geo/

### Competing Interest Statement

The authors have declared no competing interest.

